# Reproducible abnormalities of functional gradient reliably predict clinical and cognitive symptoms in schizophrenia

**DOI:** 10.1101/2020.11.24.395251

**Authors:** Meng Wang, Ang Li, Yong Liu, Hao Yan, Yuqing Sun, Ming Song, Jun Chen, Yunchun Chen, Huaning Wang, Hua Guo, Ping Wan, Luxian Lv, Yongfeng Yang, Peng Li, Lin Lu, Jun Yan, Huiling Wang, Hongxing Zhang, Dai Zhang, Tianzi Jiang, Bing Liu

## Abstract

**Background:** Schizophrenia (SZ) typically manifests heterogeneous phenotypes involving positive, negative and cognitive symptoms. However, the underlying neural mechanisms of these symptoms keep unclear. Functional gradient is a fascinating measure to characterize continuous, hierarchical organization of brain.

**Methods:** We aimed to investigate whether reproducible disruptions of functional gradient existed in SZ compared to normal controls (NC), and these abnormalities were associated with severity of clinical and cognitive symptoms in SZ. All analyses were implemented in two independent large-sample multi-site datasets (discovery dataset, 400 SZ and 336 NC; replication dataset, 279 SZ and 262 NC). First, functional gradient across cerebral cortex was calculated in each subject. Second, vertex-wise comparisons of cortical gradient between SZ and NC groups were performed to identify abnormalities in SZ. Meanwhile, reproducible and robustness analyses were implemented to validate these abnormalities. Finally, regression analyses were performed using generalized additive models to link these abnormalities to severity of clinical and cognitive symptoms in SZ.

**Results:** We found an abnormal gradient map in SZ in the discovery dataset, which was reproducible in the replication dataset. The abnormal gradient pattern was also robust when performing methodological alternatives and control analyses. Further, these reproducible abnormalities can reliably predict symptoms of clinical and cognitive domains across the two independent datasets.

**Conclusion:** These findings demonstrated that alterations in functional gradient can provide a reliable signature of SZ, characterizing the heterogenous symptoms of clinical or cognitive domains, and may be further investigated to understand the neurobiological mechanisms of these symptoms.

**Impact Statement:** In our study, using functional gradient measure and statistical learning technology and two independent multi-site case-control resting-state fMRI datasets (discovery dataset: 736 subjects; replication dataset: 541 subjects), we comprehensively investigated functional hierarchical organization in the cerebral cortex of SZ and its association with interindividual severity of symptoms. We found reproducible and robust abnormalities of functional gradient existed in SZ, which provided a reliable signature to characterize negative and general psychopathology symptoms, as well as cognitive deficits. Our findings can provide new insights to understand the neurobiological mechanisms of clinical and cognitive symptoms in SZ.

## Introduction

Schizophrenia (SZ) is commonly described as a heterogeneous mental disorder, which exhibits diverse manifestations of positive, negative and cognitive symptoms (Owen et al. 2016). Clinically, the frequency, duration and severity of symptoms usually vary significantly among individuals with SZ, and heterogeneous combinations of symptoms could be presented. Specifically, the positive symptoms are prone to remission and relapse, while negative symptoms and cognitive dysfunctions are typically persistent, which are more likely to induce lifelong disabilities (Buchanan 2007; Patel et al. 2014). However, the fundamental neurobiological mechanisms responsible for symptoms in SZ, especially the underlying neural mechanisms of these symptoms in terms of co-occurrence and interactions are yet remain unknown, which indeed hinders the development of new interventions for these symptoms.

Resting-state functional magnetic resonance imaging (rs-fMRI) has become a broadly used neuroimaging technique in psychiatric studies (Greicius 2008), and provides a non-invasive manner for investigating brain networks at macroscopic level. Plenty of rs-fMRI studies have suggested that SZ is associated with abnormalities of large-scale functional connectivity networks (van den Heuvel and Fornito 2014). However, most of these studies relied on certain parcellation strategy (Thomas Yeo et al. 2011; Wig et al. 2014; Gordon et al. 2016; Dickie et al. 2017) to define brain nodes and networks. Recently, a new approach to delineate brain organization is proposed to map functional gradients (Margulies et al. 2016; Huntenburg et al. 2018) across human cerebral cortex. Briefly, gradients refer to a series of resultant components after applying dimensionality reduction algorithms like diffusion embedding (Coifman et al. 2005) to functional connectome data from rs-fMRI. Contrary to discrete brain mapping with sharp boundaries, each gradient is regarded as a distinct axis of variation, along which cortical regions are organized continuously (Huntenburg et al. 2018), suggesting spatial transitions between cortical areas are gradual and smooth. Particularly, the first gradient is identified to explain the largest variance, which spans from unimodal sensorimotor regions to transmodal default mode network (DMN) areas (Margulies et al. 2016). Recent case-control studies demonstrate a significant disturbance along the first gradient in autism (Hong et al. 2019) and report a disrupted functional gradient of cerebellum in SZ patients (Dong et al. 2020 Mar 7), suggesting that functional gradient approach could be used to investigate neurodevelopmental disorders such as SZ. However, it is still unclear whether SZ and its clinical symptoms can be reliably characterized by aberrant functional gradient patterns across cortical regions.

In the present study, we hypothesized that reproducibly widespread or regional disruptions of functional gradient existed in SZ compared to NC, and these abnormalities could be linked to severity of clinical and cognitive symptoms. To investigate the reproducible disruptions of functional gradient in schizophrenia patients, we calculated functional gradients across cerebral cortex for each subject and examined the reliable group differences of cortical gradient in two independent large-sample case-control datasets (discovery dataset and replication dataset). Additionally, we also fully tested the robustness of gradient abnormalities in SZ with different methodological choices, such as global signal regression (Murphy and Fox 2017). Finally, we constructed regression models to predict interindividual severity of clinical and cognitive symptoms from abnormal gradients in SZ. In order to achieve interpretability, we carefully evaluated importance of gradient features and marginal effect of each feature in predictions.

## Methods

### Participants

#### Discovery dataset

The discovery dataset (n = 736, 400 SZ and 336 NC) was obtained from four Chinese hospitals: Peking University Sixth Hospital, Beijing Huilongguan Hospital, Xijing Hospital and Henan Mental Hospital, in which all subjects were scanned on 3.0T MRI scanners of Siemens. Details on participant recruitment were described in supplementary methods S1. The research protocol was approved by the Medical Research Ethics Committees of local hospitals. All participants or their guardians provided written informed consent. Subjects with any missing information in investigated clinical assessments, structural MRI or rs-fMRI were excluded (n = 63, 46 SZ and 17 NC). We censored framewise displacement (FD), and excluded subjects with mean FD greater than 0.3 mm (n = 48, 36 SZ and 12 NC). Altogether 625 individuals (318 SZ and 307 NC) remained after screening. SZ and NC groups were statistically matched in age (t = −0.70, *P* = 0.48) and gender (χ^2^ = 0.003, *P* = 0.95). Details on demographics of discovery dataset were given in supplementary table S1.

#### Replication dataset

The replication dataset included 541 individuals (279 SZ and 262 NC) who were all scanned on 3.0T MRI scanners of GE, and recruited from three hospitals in China: Renmin Hospital of Wuhan University, Zhumadian Psychiatric Hospital and Henan Mental Hospital. All clinical inclusion criterion and evaluation, data collection and quality control were identical with the discovery dataset. Written informed consent was obtained from all participants or their guardians. This study was approved by the Medical Research Ethics Committees of local hospitals. A total of 415 participants (195 SZ and 220 NC) were finally included. There were no statistically significant differences in age (t = −1.81, *P* = 0.07) and gender (χ^2^ = 0.23, *P* = 0.62) between SZ and NC groups. Details on demographics of replication dataset were given in supplementary table S1.

### Clinical assessments

Symptoms of patients with SZ were measured using Positive and Negative Syndrome Scale (PANSS) (Kay et al. 1987). Particularly, all included patients had a PANSS total score exceeded sixty and at least three of the seven positive items were scored more than four. Besides, Category Fluency Test-Animal Naming (CFT) (Strauss et al. 2006) were accomplished to assess cognitive performance in processing speed which is commonly impaired in SZ (Bokat and Goldberg 2003; Knowles et al. 2010). In CFT tasks, individuals were asked to name as many animal words as possible in one minute. A higher number of correct answers reflects a faster processing speed. Totally 497 patients (discovery dataset, 307 SZ; replication dataset, 190 SZ) finished CFT test. Details on PANSS and CFT scores were given in supplementary table S1.

### MRI acquisition and processing

Images were acquired on 3.0T Siemens scanners in the discovery dataset and obtained on 3.0T GE scanners in the replication dataset. Details on image acquisition were described in supplementary methods S2. rs-fMRI were preprocessed following steps: discarding first ten timepoints, slice timing correction, realignment, coregistration, normalisation, removing nuisances and temporal band-pass filtering (supplementary methods S3). We completed quality control by screening head motion (mean FD < 0.3 mm) and visually inspecting quality of registration and normalization. After preprocessing, we interpolated voxel-based rs-fMRI along mid-thickness surface in each individual to produce cortical vertex-based data and further resampled it to fsaverage5 space (Fischl et al. 1999). Finally, surface-based Gaussian kernel smoothing (FWHM = 2 mm) was applied.

### Functional gradient analyses

Functional gradients were calculated from rs-fMRI, referenced from prior studies (Margulies et al. 2016; Hong et al. 2019) (see supplementary methods S4 for details). We focused on analyses of the first gradient, since this dominant component explained the most variance in connectome data (supplementary figure S1) and has been suggested as a core organizing axis according with microstructure and gene expression (Huntenburg et al. 2018). In the discovery dataset, vertex-wise comparisons of gradients between SZ and NC groups were carried out by general linear models (GLMs), controlling for effects of age, gender and site (figure 1A). The significances of between-group differences were corrected for family-wise errors (*P*_FWE_ < 0.05), based on random field theory (Worsley et al. 1996) with cluster defining threshold of 0.001.

**Figure 1.**
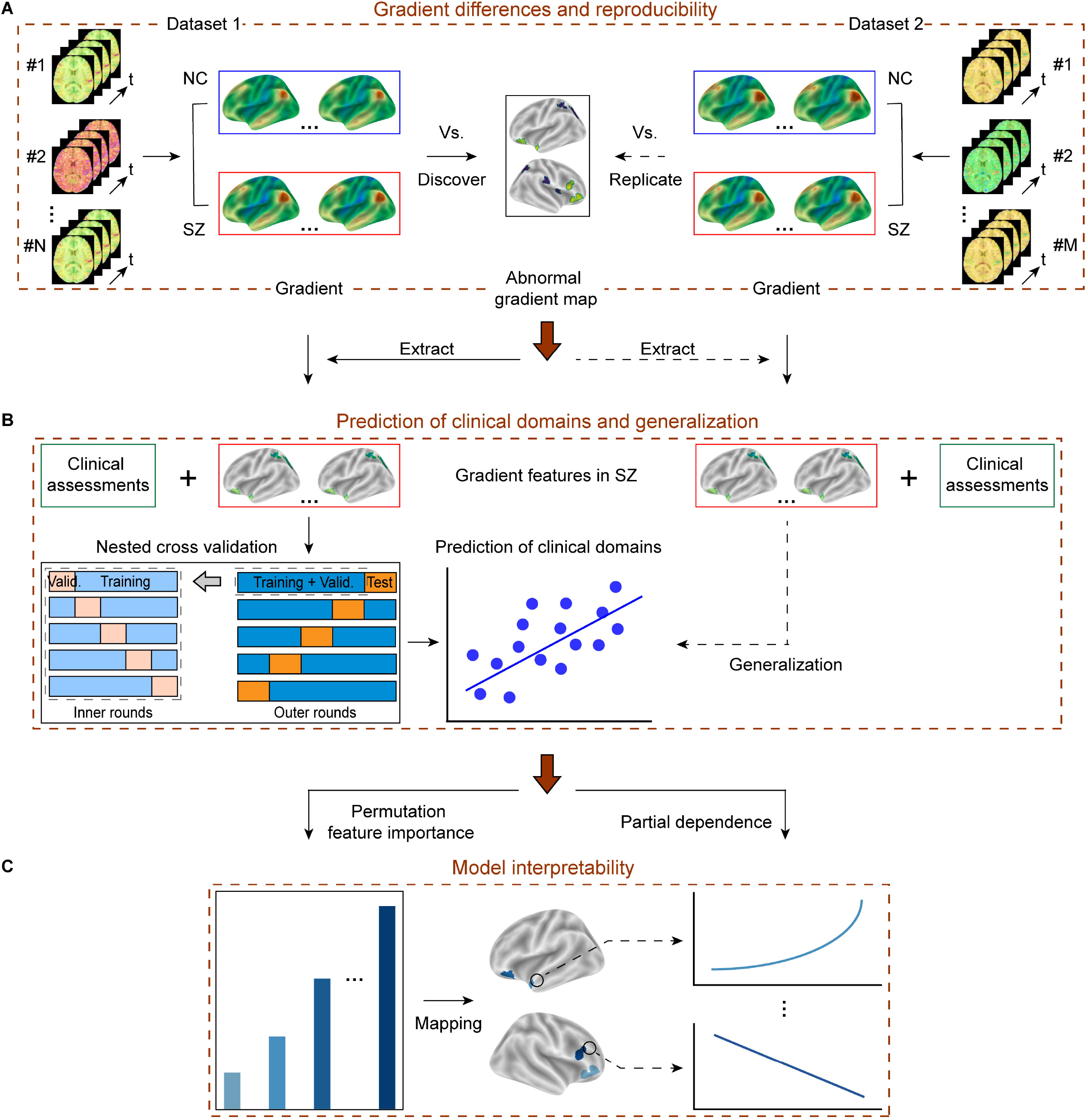
Schematic overview of identifying abnormal gradient pattern in SZ, predicting clinical characteristics and its interpretability. **A)** In the discovery dataset, between-group differences were used to establish abnormal gradient pattern in SZ. The findings were tested in an independent replication dataset. **B**) Gradient values within abnormal clusters were used as features to predict clinical assessments. Regression models were built, trained and tested in the discovery dataset, and further tested the generalization performance in the replication dataset. **C**) Evaluated feature importance and marginal effects.

### Reproducible and robustness analyses of gradient findings

To validate the reproducibility of abnormal gradient patterns, in the replication dataset, we extracted and averaged gradient values among all identified clusters (more than 30 connected vertices, *P*_FWE_ < 0.05 in the discovery dataset). Similarly, GLMs were applied to conduct cluster-wise comparisons between SZ and NC groups, with age, gender and site controlled as covariates (figure 1A). False discovery rate (FDR) correction was applied at several thresholds (0.05, 0.005 and 0.001).

Several alternative methodological choices were tested to further ensure the robustness of our results: i) controlling mean framewise displacement (Van Dijk et al. 2012). ii) performing global signal regression (GSR) (Murphy and Fox 2017). iii) different thresholds when constructing individualized functional connectivity network. More details are provided in supplementary method S4.

### Prediction of symptom severity and cognitive performance

#### Generalized additive models (GAMs)

Regression analyses were conducted to investigate whether identified abnormal gradient patterns were associated with clinical symptoms and cognitive ability in processing speed in SZ (figure 1B). PANSS total and subscale (positive, negative and general psychopathology) scores were used to evaluate symptom severity, processing speed was measured by CFT scores. We built predictive models under the framework of GAMs (Hastie and Tibshirani 1986; Hastie and Tibshirani 1995), which is a powerful and yet simple technique combining advantages of interpretability, flexibility/automation and regularization together. See supplementary methods S6 for details.

#### Model training, testing and generalization

We generated an abnormal gradient map by combining all significant clusters derived from between-group comparisons in the discovery dataset. Mean gradient values extracted from clusters were used as features, clinical assessments were targets. We trained models via nested 5-fold cross-validation (CV) (Varoquaux et al. 2017) in the discovery dataset (figure 1B), which can effectively avoid biases in model selection (Cawley and Talbot 2010). See supplementary methods S6 and figure S5 for details. To further evaluate generalization performance, we selected the predictive model with best testing performance, and validated in the replication dataset after retrained on the whole discovery dataset. To quantify the quality of predictions, Pearson correlation and mean absolute error (MAE) were calculated between observed and predicted scores. A great number of permutation tests (100,000 times) were completed by randomly shuffling targets to acquire the significance levels of predictions. Similarly, all gradient features were adjusted for age, gender and site.

#### Model interpretability

Permutation feature importance (Molnar 2020) was used to evaluate importance of each feature in predictive models (figure 1C). Firstly, variance inflation factor (VIF) (James et al. 2013) was calculated to detect multicollinearity among features. Then, we repeated nested 5-fold CV with one feature randomly shuffled, and calculated percentage reduction in performance to characterize its importance in prediction (supplementary methods S7). To evaluate dependences between features and target, we used partial dependence plot (PDP) (Molnar 2020) to show marginal effects of features had on predicted target (figure 1C). Particularly, PDP describes whether the relationship between a feature and target is linear, monotonic or complicated.

#### Alternative models analyses

We also built several other commonly used regression models to test whether our gradient features can predict clinical targets by other algorithms. Nested 5-fold CV or single 5-fold CV was used depending on whether the model had hyperparameters to tune. All other procedures and performance evaluation were the same with GAMs.

## Results

### Spatial similarity of gradient between schizophrenia and normal controls

In both discovery and replication datasets, the first gradient explained 25% connectome variance, and there were no differences in amount of explained variance between SZ and NC groups (FDR adjusted *P* > 0.05; supplementary figure S1). The first gradient presented gradual connectivity variations along an embedding axis, which anchored the unimodal primary sensory systems at one end and the integrative transmodal DMN regions at the other end (figure 2A and figure 2C), resemble recent findings (Margulies et al. 2016; Hong et al. 2019; Dong et al. 2020 Mar 8). This gradient pattern was highly consistent across SZ and NC groups in spatial (*r* = 0.98; figure 2B and figure 2D) and reflects a well replication for a previous study establishing this gradient in a Caucasian population (Margulies et al. 2016) (*r* = 0.86; supplementary figure S2).

**Figure 2.**
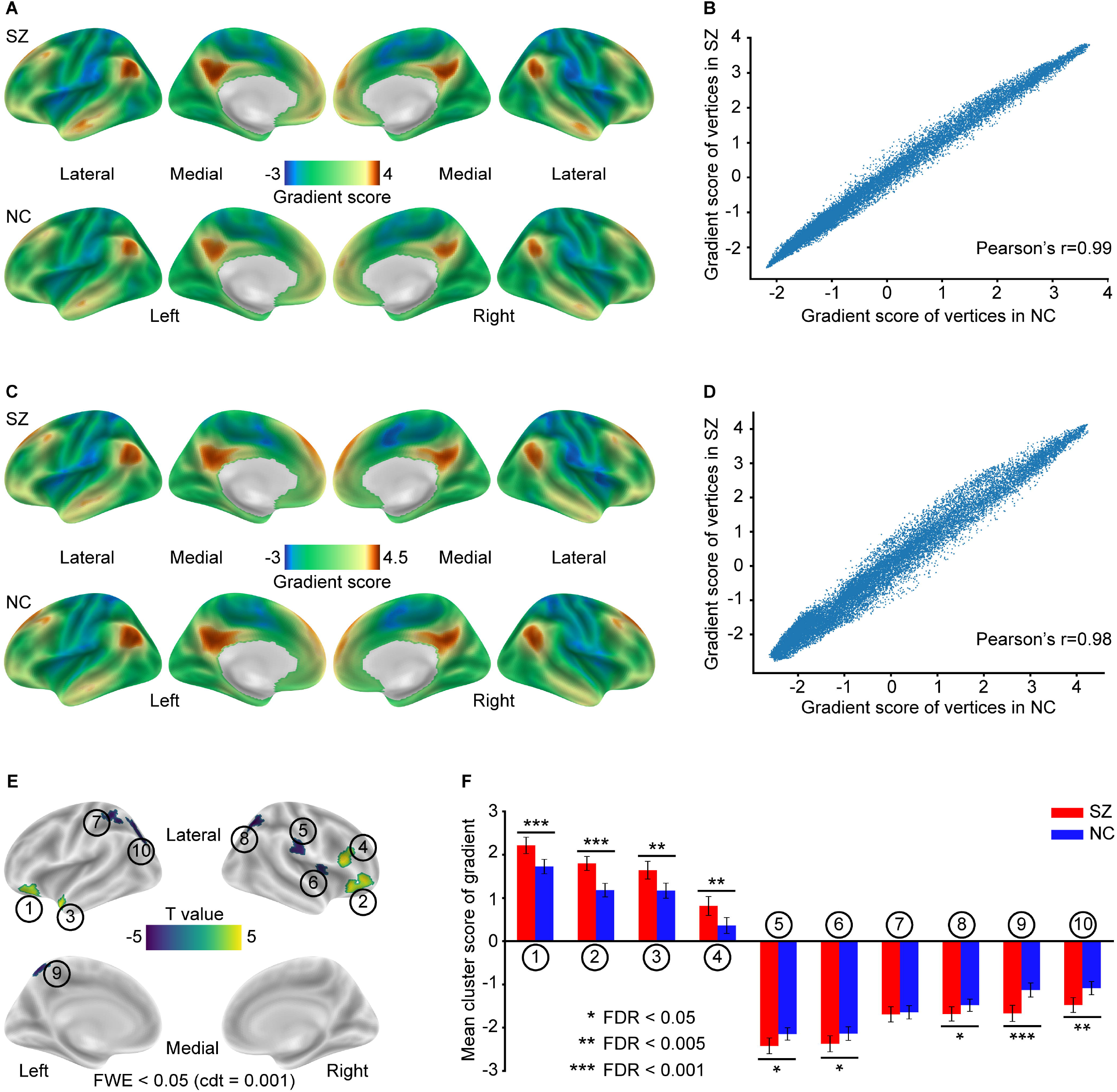
Gradient mapping on cortical surface. Gradient patterns in SZ and NC groups in the discovery (**A**) and replication (**C**) datasets. Spatial correlation of gradient between groups in the discovery (**B**) and replication (**D**) datasets. **E**) In the discovery dataset, ten significant clusters of between-group differences in gradient were shown with circle numbers and t-statistic values. **F**) In the replication dataset, the effects of these ten clusters can be replicated (except for cluster 7). Error bars indicated 95% confidence interval.

### Reproducible and robust abnormalities of gradient in schizophrenia

In the discovery dataset, ten significant clusters were identified when performing vertex-wise comparisons of gradient between SZ and NC groups (*P*_FWE_ < 0.05, cdt = 0.001; figure 2E). Compared with NC, gradient in SZ increased in positive range, centered on transmodal regions including bilateral orbitofrontal cortex (OFC; cluster 1 and cluster 2), left temporal pole (TPO; cluster 3), and right dorsolateral prefrontal cortex (DLPFC; cluster 4). Decreased regions in SZ were located at negative range, including right supramarginal gyrus (SMG; cluster 5), right insula (INS; cluster 6), left inferior parietal (IPL; cluster 7), bilateral superior parietal gyrus (SPG; cluster 8 and cluster 10), and left precuneus (PCUN; cluster 9). Detailed cluster information were given in supplementary table S2. Importantly, these comparing results could be generalized to replication dataset (figure 2F). Particularly, between-group differences of mean gradient scores in the ten clusters (except cluster 7) can be replicated (FDR adjusted *P* < 0.05), and six clusters remained significant after FDR correction at a stringent level of 0.005. Detailed replicated results were given in supplementary table S3. Moreover, robustness analyses indicated that our results were of highly consistency. Specifically, significant clusters can be well replicated when i) regressing out mean FD (supplementary figure S3A); ii) using GSR-processed rs-fMRI (supplementary figure S3B); iii) thresholding connectivity matrix at different proportions from 5% to 25% (supplementary figure S4). These validation findings suggested that disrupted gradient pattern in SZ was reliable and reproducible.

### Abnormal gradients predict clinical and cognitive domains in schizophrenia

Having identified reproducible alterations of gradient, we tested if these abnormalities can characterize symptom severity and cognitive deficits in SZ. Regression analyses were performed by GAMs, model training and testing were implemented through nested 5-fold CV in the discovery dataset, generalization performance was further evaluated in the replication dataset. We found cluster-based gradient features can significantly predict PANSS total scores (figure 3A, discovery dataset: mean *r* = 0.28, *P* < 1 × 10^-5^, MAE =10.2; figure 3B, replication dataset: *r* = 0.26, *P* = 4 × 10^-5^, MAE = 12.87) and subscale scores of negative (figure 3C, discovery dataset: mean *r* = 0.22, *P* = 2.5 × 10^-4^, MAE = 5.35; figure 3D, replication dataset: *r* = 0.22, *P* = 7.2 × 10^-4^, MAE = 5.77) and general psychopathology (figure 3E, discovery dataset: mean *r* = 0.26, *P* = 5 × 10^-5^, MAE = 7.02; figure 3F, replication dataset: *r* = 0.23, *P* = 6.2 × 10^-4^, MAE = 7.20), as well as CFT scores (figure 3G, discovery dataset: mean *r* = 0.25, *P* = 1 × 10^-5^, MAE = 5.28; figure 3H, replication dataset: *r* = 0.17, *P* = 0.009, MAE = 7.05). Interestingly, no effect can be observed in predicting PANSS positive scale (*P* > 0.05; supplementary table S4). Furthermore, we found domains of PANSS total and subscales of negative and general psychopathology can also be predicted by other algorithms (but not by simple linear algorithms; supplementary table S5), indicating that predictive ability of functional gradients was not restricted to a particular model.

**Figure 3.**
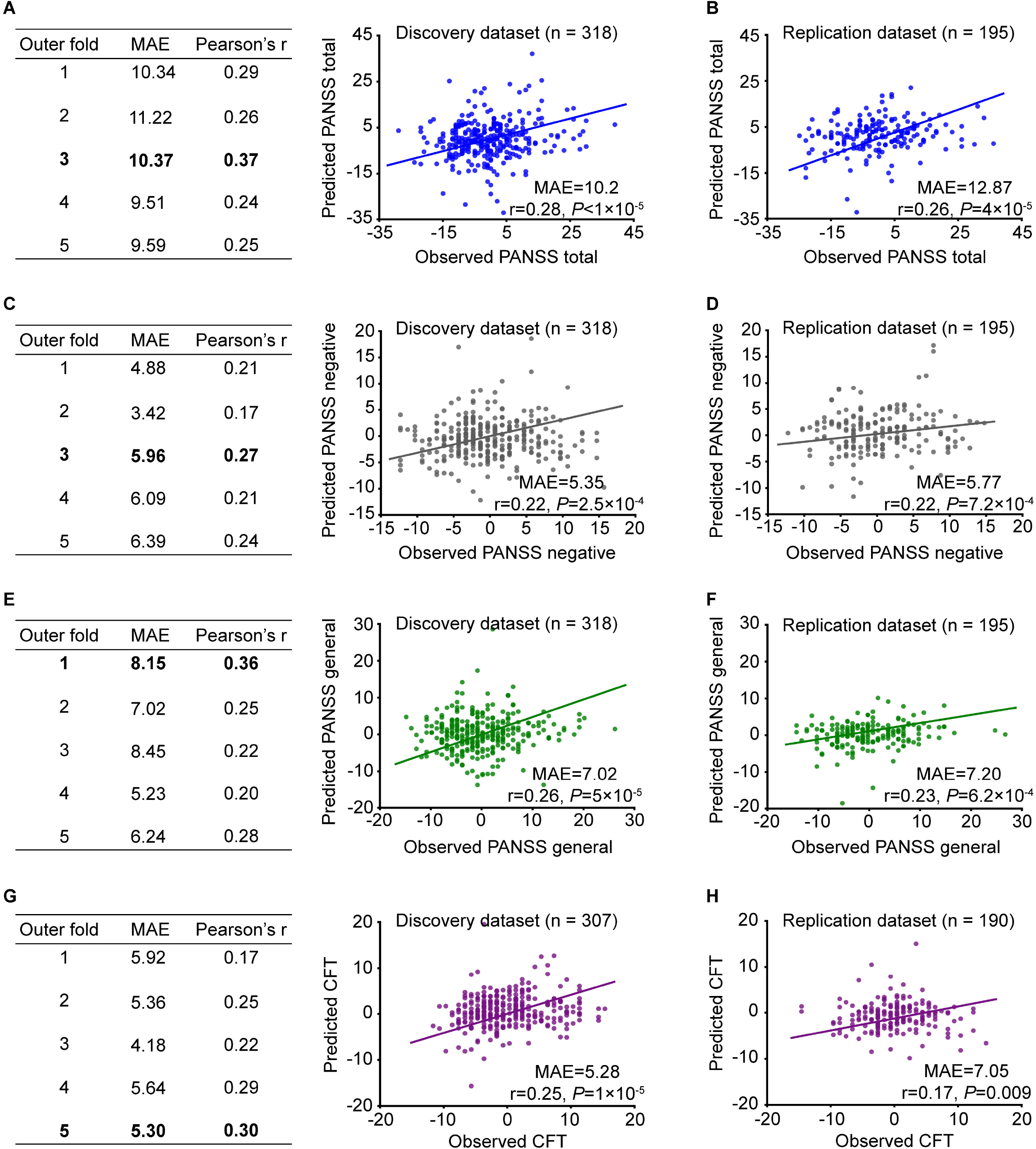
Abnormal gradients predicted clinical domains in SZ in the discovery and replication datasets. Significant prediction models can be achieved to characterize PANSS total scores (**A** and **B**) and subscale scores of negative (**C** and **D**) and general psychopathology (**E** and **F**) as well as CFT scores (**G** and **H**). Testing results of nested 5-fold CV were provided in left table. Scatter plots shown mean testing performance in the discovery dataset and generalization performance in the replication dataset.

### Gradient importance and marginal effect in predictions

We used permutation feature importance and PDP to characterize importance and marginal effects of gradient features. VIF results (all less than 5; supplementary table S6) indicated features were of minor multicollinearity. We found gradients in OFC, TPO and DLPFC were reliably higher importance in predictions of PANSS domains (figure 4B, figure 4C and figure 4D), and especially gradient in DLPFC was the most important feature. However, when predicting CFT domain, these four features were less important, and gradient in SPG became the most significant one (figure 4E). Detailed information was provided in supplementary table S7.

**Figure 4.**
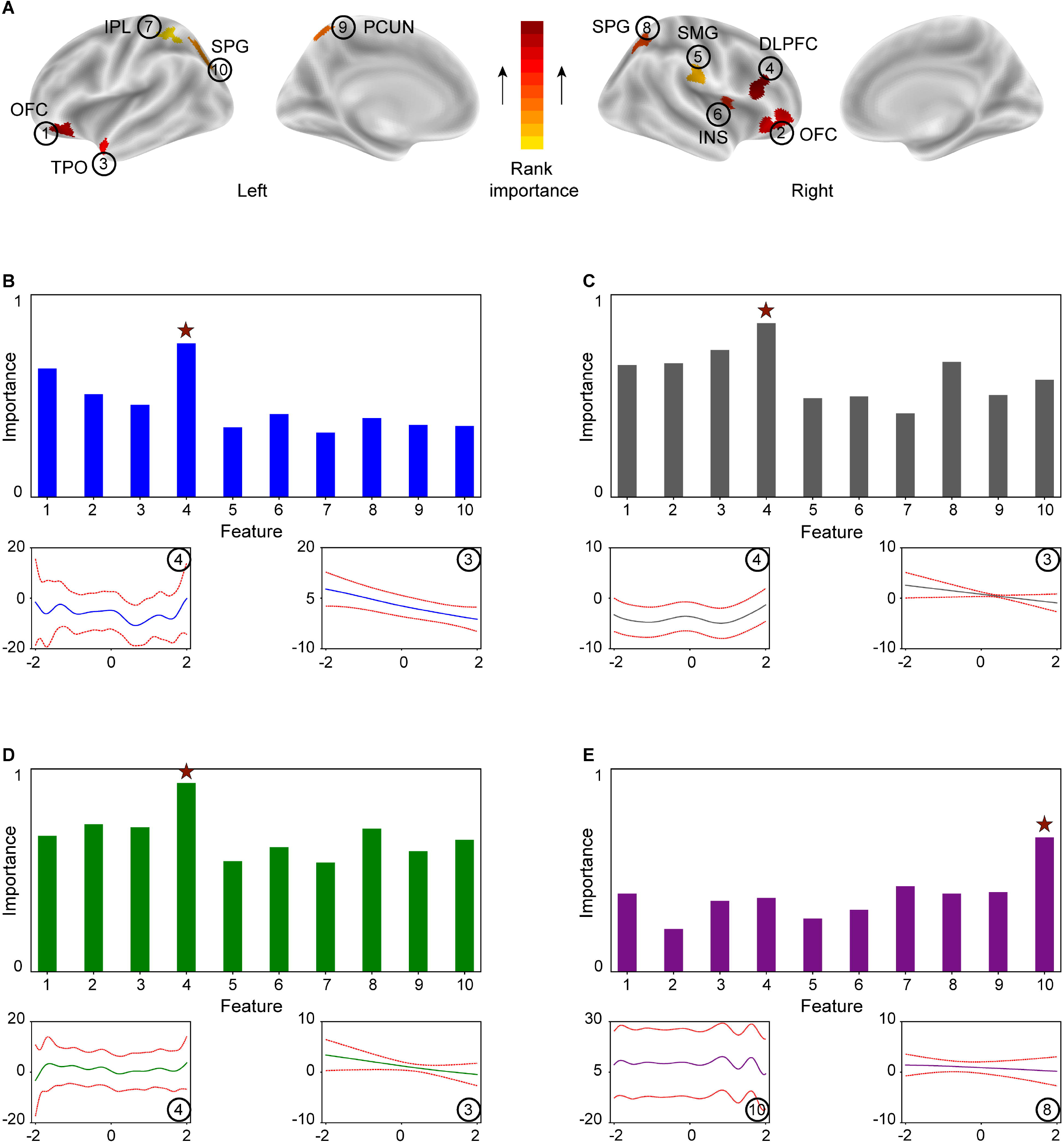
Contributions of gradient features in prediction models. **A**) Ranking importance of gradient features when predicting PANSS total scores. Specifically, feature importance and PDPs of feature 4 and feature 3 were shown when **B**) predicting PANSS total scores; **C**) subscale scores of negative and **D**) general psychopathology. **E**) Feature importance and PDPs of feature 10 and feature 8 when predicting CFT scores. The two red lines in PDPs were 95% confidence interval. Features were labelled with circle numbers. The most important predicted feature was marked with red star in bar chats.

When examining dependences between features and target, marginal effects were diverse. Clearly, relationships between TPO feature and three PANSS domains were linear. Nevertheless, more complex relationships were found between DLPFC feature and three PANSS domains (figure 4B, figure 4C and figure 4D). Similarly, relationships between features and CFT domain also shown linearity and complexity (figure 4E). Complete PDPs were provided in supplementary figure S6. This result may explain that using simple linear algorithms cannot reliably predict clinical domains (supplementary table S5).

## Discussion

Identification of brain network substrates of psychiatric symptoms has long been a primary and challenging goal in SZ studies. In our study, we comprehensively studied functional gradients, which is supplementary to traditional discrete mapping of brain networks, to investigate whether this new approach can help explaining the neural mechanisms underlying clinical or cognitive symptoms in SZ. Specifically, we established that the first gradients were highly spatially correlated between SZ and NC groups, expanding from unimodal somatosensory regions to transmodal DMN areas. We carefully revealed and validated the abnormal gradient pattern in SZ at cluster levels. Through regression analyses, we found these abnormalities of gradient can significantly predict clinical symptoms and cognitive deficits in SZ, and the contributions of features were fully evaluated. The independent validation strategy, methodological alternatives and control analyses demonstrated the reliability and reproducibility of our results, which are indispensable issues in neuroimaging studies (Button et al. 2013; Zuo et al. 2019).

One end of the first gradient was located at primary somatosensory/motor areas, which are directly related with low-level brain functions such as early sensory processing (Borich et al. 2015). Transmodal DMN regions, which are known to be engaged in a variety of higher-level cognitions (Anticevic et al. 2012) were situated at the other end. Therefore, the first gradient, varied between the two opposite ends on cortex, reflected a hierarchy of brain functions, integrating both low- and higher-levels, which are suggested to be impaired in SZ (Javitt 2009; Fioravanti et al. 2012). Although principle functional gradient patterns in SZ and NC groups were highly consistent and explained similar variance, we revealed several significant regional alternations in SZ. Specifically, gradients were increased in bilateral OFC, left TPO and right DLPFC, and decreased in right SMG, right INS, left IPL, bilateral SPG and left PCUN in SZ. Importantly, these findings were well reproduced across two independent datasets, and were robust when controlling for confounding variables such as mean framewise displacement (Power et al. 2012), and considering effects of different methodological choices such as whether or not performed GSR (Murphy and Fox 2017) in rs-fMRI data, and multiple thresholding proportions in connectivity matrix. Among these regions, abnormalities in OFC, DLPFC, TPO, PCUN and SPG have larger effect sizes. As we have known, OFC, DLPFC and TPO are generally involved in a wide variety of higher order cognitive processes such as decision making, cognitive flexibility, executive functions and semantic memory (Krawczyk 2002; Rolls 2004; Olson et al. 2007). PCUN and SPG are typically related to perception and sensation (Cavanna and Trimble 2006). Meanwhile, a number of previous rs-fMRI studies have also discovered abnormal functional connectivity involving most of these regions (Mitchell et al. 2001; Karbasforoushan and Woodward 2012). However, our identified abnormal brain regions were not fully consistent with a recent gradient study on SZ (Dong et al. 2020 Mar 8). This inconsistent might be ascribed to differences in sample size (Button et al. 2013), clinical heterogeneity, image processing, data quality control and statistical procedures such as multiple testing methods and thresholds. In brief, we identified robust and reproducible abnormalities of gradient in SZ compared to NC.

To investigate associations between abnormal gradients and clinical symptoms in SZ, we performed regression analyses using generalized additive models (GAMs) (Hastie and Tibshirani 1986; Hastie and Tibshirani 1995). Particularly, selection of GAMs was the result of overall considerations. First, GAMs have good advantages in both transparency and model capacity (Bzdok and Ioannidis 2019), meaning we can estimate complex nonlinear relationships while maintaining model simplicity. Second, the number of gradient features was relatively small (only 10), compared to typical neuroimaging studies, and thus can avoid the parameter-heavy problem in GAMs.

Third, gradient features were not multicollinear, quantified by variance inflation factor (VIF) (James et al. 2013), since GAMs are not immune to multicollinearity. Results proved that abnormal gradients can reliably predicted PANSS total, subscales of negative and general psychopathology as well as CFT scores. Interestingly, we found that PANSS positive scale cannot be predicted, suggesting a different neurobiological mechanism in this domain. Importantly, we found that the prediction can be well generalized to an independent replication dataset, establishing the potential of external validation. Collectively, our results suggested these alterations in functional gradient may reflect a reliable pathological signature characterizing negative and general psychopathology symptoms, and CFT cognitive domain in SZ, rather than positive symptom domain.

When evaluating importance of gradient features, we consistently found gradient in DLPFC was the most important feature in predictions of PANSS total, subscales of negative and general psychopathology. Actually, several studies have reported that aberrant DLPFC-related connectivity were linked to symptoms in SZ. For example, a recent study indicated that connectivity breakdown between DLPFC and cerebellum is associated with negative symptom severity (Brady et al. 2019); reduced connectivity between DLPFC and dorsal parietal cortex is also reported to be linked with dysfunctions in executive function, working memory and attention (Barch and Ceaser 2012). Besides, secondary important features such as gradient in OFC and TPO also had great contributions on predictions of PANSS domains. Indeed, studies have suggested that OFC-related connectivity is associated with negative symptom severity (Shukla et al. 2018 Dec 21) (right OFC and striatum), and general psychopathology score (left OFC and right precuneus) in first-episode schizophrenia (Li et al. 2016 Jul 21). Although gradient in IPL was least important relative to other features, it was still useful in predictions of PANSS domains, since performance decreased at a non-negligible level (at least 17%) when it was permuted. Meanwhile, functional differences in IPL were also observed in functions of sensory integration and executive functions in SZ (Torrey 2007). As for dependences between gradient features and clinical scores, most of gradient features had complex nonlinear marginal effects. These can also be verified that all linear models cannot predict clinical domains.

Despite reliable associations between abnormalities of gradient and symptoms in SZ have been revealed, there is a major limitation to be considered and addressed in future work. Our cross-sectional design may restrict the abilities to validate the causal relationships between neuroimaging and clinical assessments. Therefore, longitudinal data should be included to further test how gradients changed with ongoing symptoms, and whether gradients can still reliably predict clinical symptoms in follow-up samples.

## Conclusion

Our work revealed significant, robust and reproducible alterations in functional gradients of SZ across two independent datasets, and found these abnormalities of gradient can reliably predict PANSS total, subscales of negative and general psychopathology as well as CFT scores. Our findings provided a novel insight to understand the functional abnormalities and diverse symptoms in SZ.

## Acknowledgements

We thank all the patients and controls engaged in this study, as well as the clinical staff who facilitated their involvement.

## Author contributions

M.W. and B.L. performed data analyses and prepared the manuscript. A.L. made substantial contributions to the manuscript and provided critical comments. Y.L., Y.S., M.S., and T.J. participated in discussions of the results. H.Y, J.C., Y.C., H.W., H.G., P.W., L.L., Y.Y., P.L., L.L., J.Y., H.W., H.Z. and D.Z. contributed to data acquisition.

## Conflict of Interest

The authors declare no competing interests.

## Funding Statement

This work was supported by the National Key Basic Research and Development Program (973) (Grant 2011CB707800), the Strategic Priority Research Program of Chinese Academy of Science (Grant No. XDB32020200), and the Natural Science Foundation of China (Grant numbers 81771451).

## Notes

### Competing Interest Statement

The authors have declared no competing interest.

