## Supplementary Material for "Reproducible abnormalities of functional gradient reliably predict clinical and cognitive symptoms in schizophrenia"

**Methods S1.** Participant recruitment

All individuals with SZ were diagnosed consensually by two qualified psychiatrists utilizing the Structured Clinical interview for DSM-IV Axis I Disorders (SCID-I/P, Patient Edition). None of the included patients had any history of other DSM-IV Axis I disorders, neurological disorders, cognitive disabilities, serious physical diseases, severe head trauma, substance abuse or dependence and electroconvulsive therapy within the last 6 months. All healthy subjects were assessed clinically by SCID-I/NP (Non-patient Edition), who and their families (first- and second-degree relatives) had no history of any mental disorders.

**Methods S2.** Image acquisition

In the discovery dataset, images were acquired on 3.0T Siemens scanners: TrioTim in Peking University Sixth Hospital (PKU6), Beijing Huilongguan Hospital (HLG) and

Xijing Hospital (XIAN); Verio in Henan Mental Hospital (XX-Siemens). For rs-fMRI, 2D echo planar imaging (EPI) sequence was used with parameters: repetition time (TR) = 2000 ms; echo time (TE) = 30 ms; flip angle (FA) = 90$^{o}$; field of view (FOV) = 220 × 220 mm$^2$; matrix size = 64 × 64; voxel size = 3.4375 × 3.4375 × 4.6 mm$^3$; 240 volumes, 33 slices. For T1-weighted images (T1w), 3D-MPRAGE sequence was performed with parameters: TR = 2530 ms; TE = 3.5 ms (PKU6, HLG and XIAN) or 2.43 ms (XX-Siemens); FA = 7$^{o}$; inversion time (TI) = 1100 ms; voxel size = 1 × 1 × 1 mm$^3$; matrix size = 256 × 256 × 192.

In the replication dataset, images were acquired on 3.0T GE scanners: Signa HDxt in Renmin Hospital of Wuhan University (WUHAN), Zhumadian Psychiatric Hospital (ZMD) and Henan Mental Hospital (XX-GE). For rs-fMRI, 2D-EPI sequence was used with parameters: TR = 2000 ms; TE = 30 ms; FA = 90$^{o}$; FOV = 220 × 220 mm$^2$; matrix size = 64 × 64; voxel size = 3.4375 × 3.4375 × 4.6 mm$^3$; 240 (WUHAN and XX-GE) or 180 (ZMD) volumes, 32 (WUHAN) or 33 (ZMD and XX-GE) slices. For T1w, 3D-GRE sequence was performed with parameters: TR = 7.8 ms (WUHAN and XX-GE) or 6.8 ms (ZMD); TE = 3 ms (WUHAN and XX-GE) or 2.5 ms (ZMD); FA = 7$^{o}$; TI = 1100 ms; voxel size = 1 × 1 × 1 mm$^3$; matrix size = 256 × 256 × 188.

**Methods S3.** rs-fMRI preprocessing

All rs-fMRI data from both discovery and replication datasets were preprocessed with identical, standardized procedures based on BRANT (Xu et al. 2018) (https://github.com/kbxu/brant), an open source MATLAB-based fMRI toolkit integrated with built-in scripts from SPM (Eickhoff et al. 2005). Briefly, the following successive stages were contained: removing the initial ten timepoints, slice timing correction, within-subject realignment, coregistration from T1w to EPI mean image by rigid-body transformation, spatial normalisation of EPI images to standard space Montreal Neurological Institute (MNI) using the segmentations of T1w, resampling the normalised EPI images to $3 \times 3 \times 3$ mm$^3$, removing nuisances by regressing out linear trend, mean time series from white matter and cerebrospinal fluid as well as estimated head-motion parameters, and temporal band-pass filtering at 0.01 - 0.08 HZ.

**Methods S4.** Calculation of functional gradient

For each individual in the discovery and replication datasets, vertex-wise functional gradients across entire cortical regions were calculated based on resting-state functional connectome data, referenced from previous studies (Margulies et al. 2016; Hong et al. 2019). Specifically, using rs-fMRI time series on cortical surface of each subject, we first constructed functional connectivity matrix (rsFC, 20,484 × 20,484 entries) based on Pearson correlation. We then z-transformed and thresholded rsFC by reserving the values of top 10% connections in each row, while zeroing all others and negative connections, and further created affinity matrix by calculating cosine distance between all pairs of rows and subtracting it by one. The resulted affinity matrix, in which values ranged from zero to one, was symmetric and captured the similarity of connectome profiles between vertices (Margulies et al. 2016). After that, we performed dimensionality reduction on the affinity matrix by applying a nonlinear manifold learning algorithm, diffusion maps (Coifman et al. 2005), which was determined by one single parameter α. We followed the choices by prior studies (Margulies et al. 2016; Guell et al. 2018; Hong et al. 2019), and assigned α to 0.5, which was demonstrated well-suited for functional connectivity data (Margulies et al. 2016). This algorithm eventually estimated several principal components in low-dimensional embedding space, referred to as functional gradients, each of which (1 × 20,484 entries) explained a ratio of connectome variance. Finally, in each dataset, we separately created a group-averaged template based on obtained gradients in all individuals (both SZ and NC groups were engaged), and performed orthonormal alignment from gradients of each subject to the template via Procrustes analysis (Langs et al. 2015).

**Methods S5.** Robustness analyses of between-group differences in gradient

Several alternative analyses were performed in the discovery dataset to demonstrate robustness of abnormal gradient pattern in SZ.

1) Considering that head motion is a confounding factor had significant, systematic impacts on rs-fMRI connectivity measures (Van Dijk et al. 2012), we repeated between-group comparisons while additionally including mean framewise displacement (Power et al. 2012) as a covariate in the GLM models.

2) Whether including global signal regression (GSR) in the preprocessing of rs-fMRI has long been a controversial issue, and does not reached a consensus (Murphy and Fox 2017). In our main analysis, we did not perform GSR for rs-fMRI. Nevertheless, to

test the effects of GSR on gradients, we reprocessed rs-fMRI in each individual by including GSR, and remained all other procedures the same. We then repeated gradient analyses using GSR-processed data.

3) To examine effects of different proportional thresholds in connectivity matrix, we additionally selected four thresholds (5%, 15%, 20% and 25%), and repeated gradient analyses respectively.

**Methods S6.** Building and training generalized additive models (GAMs)

For simplicity, linear GAMs were modeled which extended general linear models by allowing nonlinear functions of predictor variables (feature functions) while maintaining additivity. The feature functions are generally smooth, and are able to capture nonlinear relationships between each predictor variable and response variable. In our analysis, penalized B splines (Eilers and Marx 1996) were used as feature functions, which allowed us to control the smoothness by a penalty parameter to prevent overfitting.

Model training was completed by a nested 5-fold cross-validation (CV) (Varoquaux et al. 2017) design. The nested 5-fold CV structure contains inner and outer procedures, which are used for tuning penalty (smoothing) parameters of feature functions and testing performance respectively. Besides, randomized search strategy (Bergstra and Bengio) (10,000 times) was applied to estimate smoothing parameters, considering the high-dimensional search-spaces.

**Methods S7.** Gradient feature importance

We first calculated variance inflation factor (VIF) (James et al. 2013) to detect multicollinearity among features, considering models could still have access to permuted feature through its correlated features and thus leading to misleading values of importance. We then repeated nested 5-fold CV with each single feature randomly shuffled, and calculated the percentage reduction of Pearson correlation between predicted and observed value in the discovery dataset. This procedure can break the correspondence between features and target, and thus drops of model performance can indicate how much the model depends on the feature. Each feature was separately permuted 100 times, and we chose the median value of percentage reductions in Pearson correlations to quantify its importance.

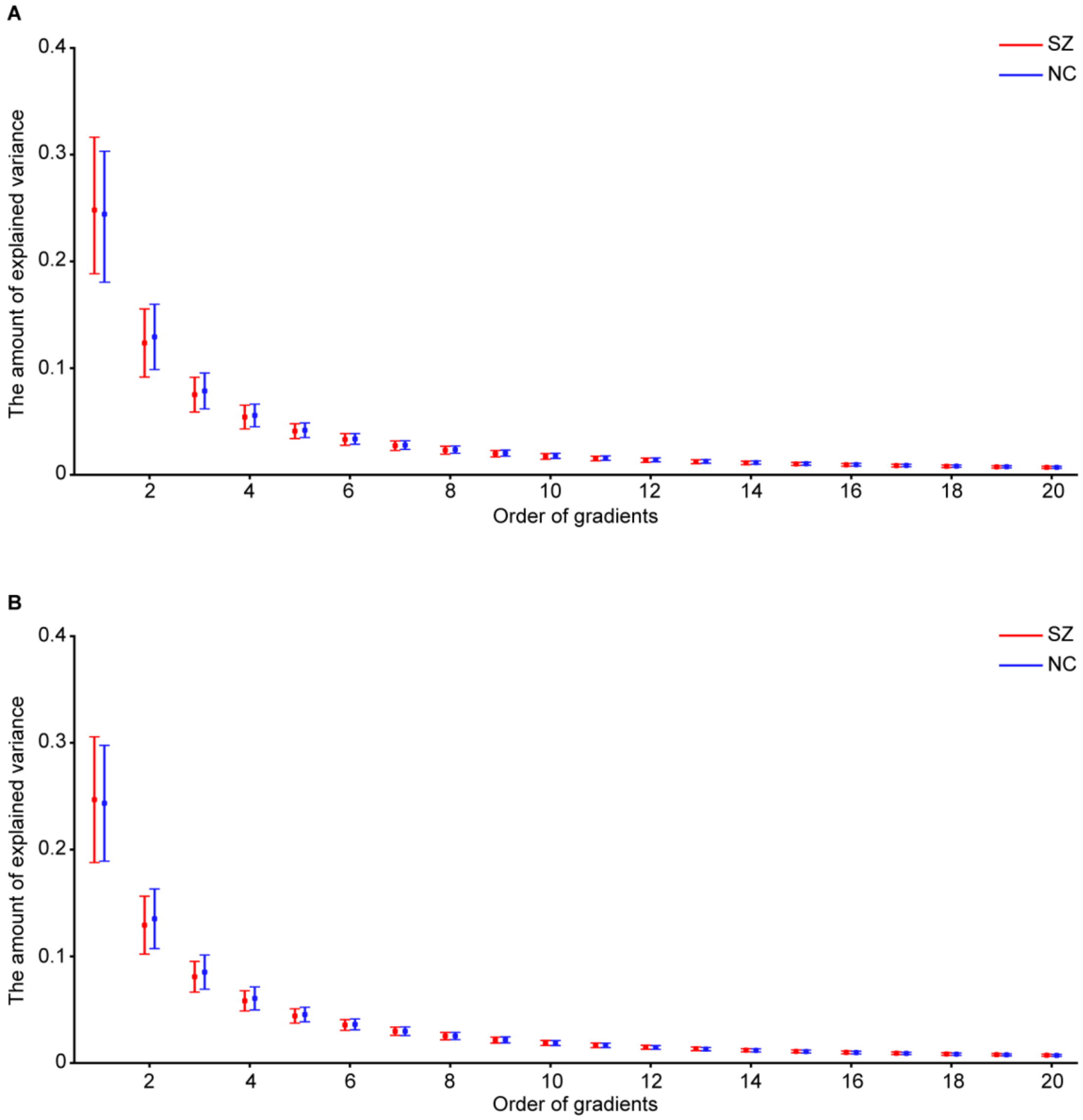


**Figure S1.** Explained variance of the first 20 gradients in SZ and NC groups in the discovery (**A**) and replication (**B**) datasets. In each dataset, the first gradient explained the most amount of variance, approximately 0.25, in both SZ and NC groups. There were also no statistically significant between-group differences in the amount of explained variance across gradients. Error bars represented one standard deviation.

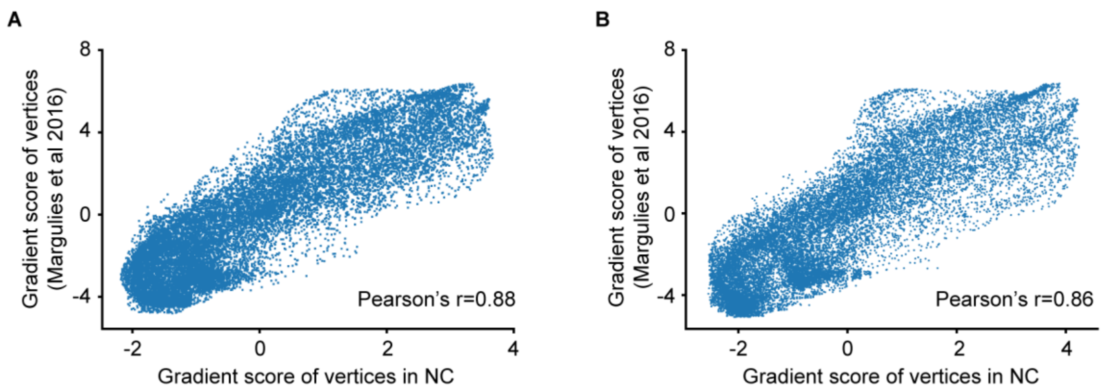


**Figure S2.** Spatial correlation of gradient score in vertices between a previous study of healthy subjects and NC group in the discovery (**A**) and replication (**B**) datasets.

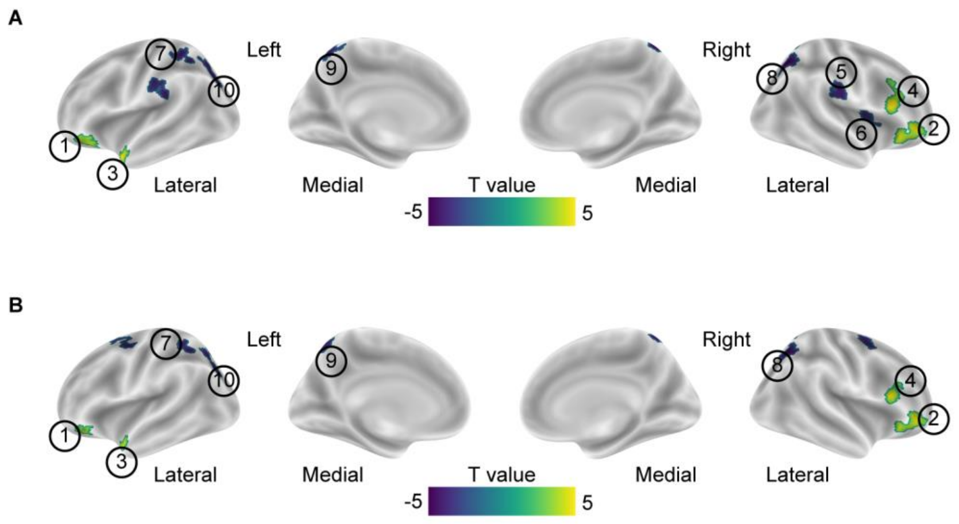


**Figure S3.** Consistency of between-group differences in gradient while additionally regressing out mean framewise displacement (**A**) and using global signal regression (GSR) processed rs-fMRI data (**B**) in the discovery dataset. Statistical procedures were the same as for figure 2E. T maps were shown, and the significant clusters labeled with circle numbers were corresponded to those in figure 2E.

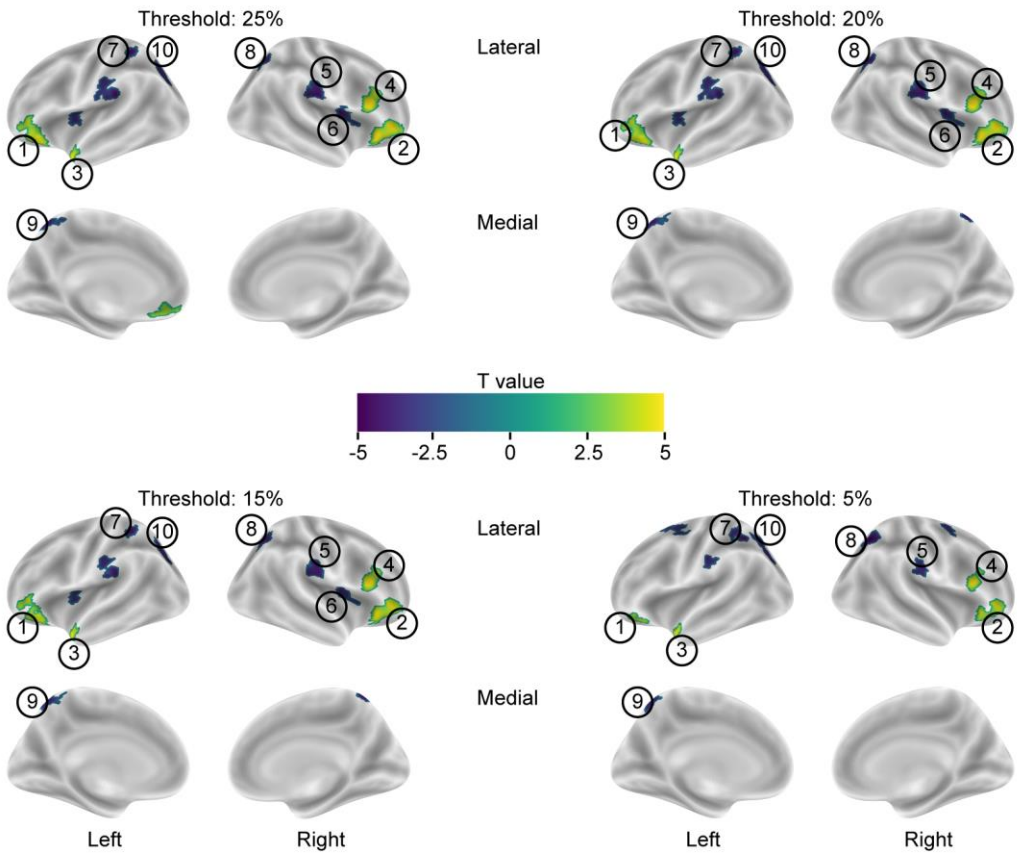


**Figure S4.** Consistency of between-group differences in gradient when applying different thresholds (5% to 25%) in functional connectivity matrix. Statistical procedures were the same as for figure 2E. T maps were shown, and the significant clusters labeled with circle numbers were corresponded to those in figure 2E.

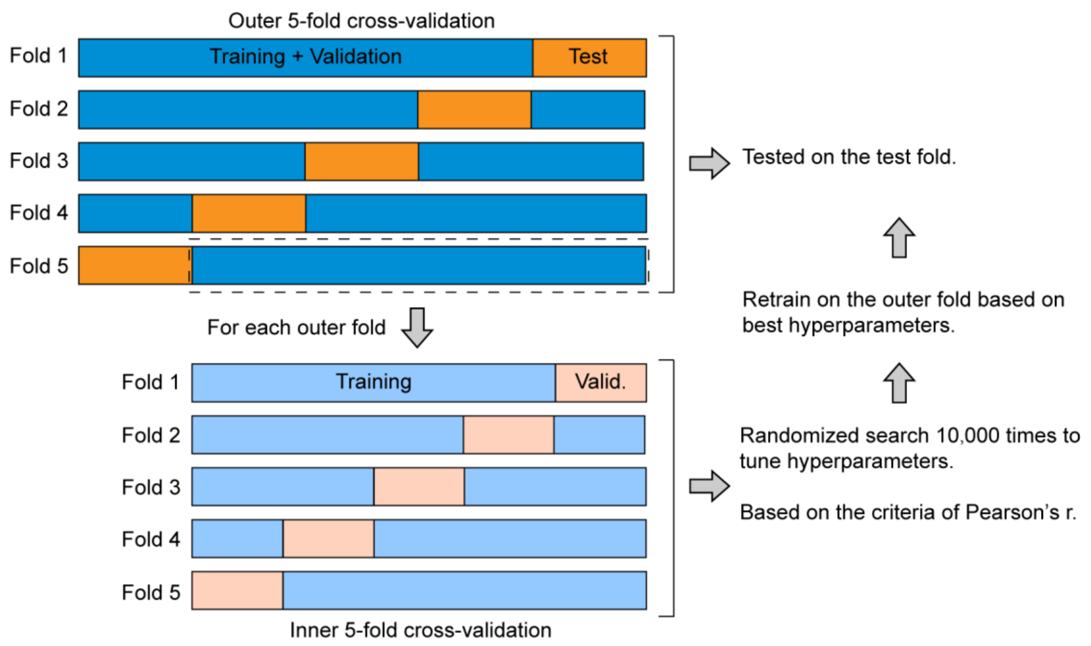


**Figure S5.** The process of nested 5-fold cross-validation (CV) in the discovery dataset.

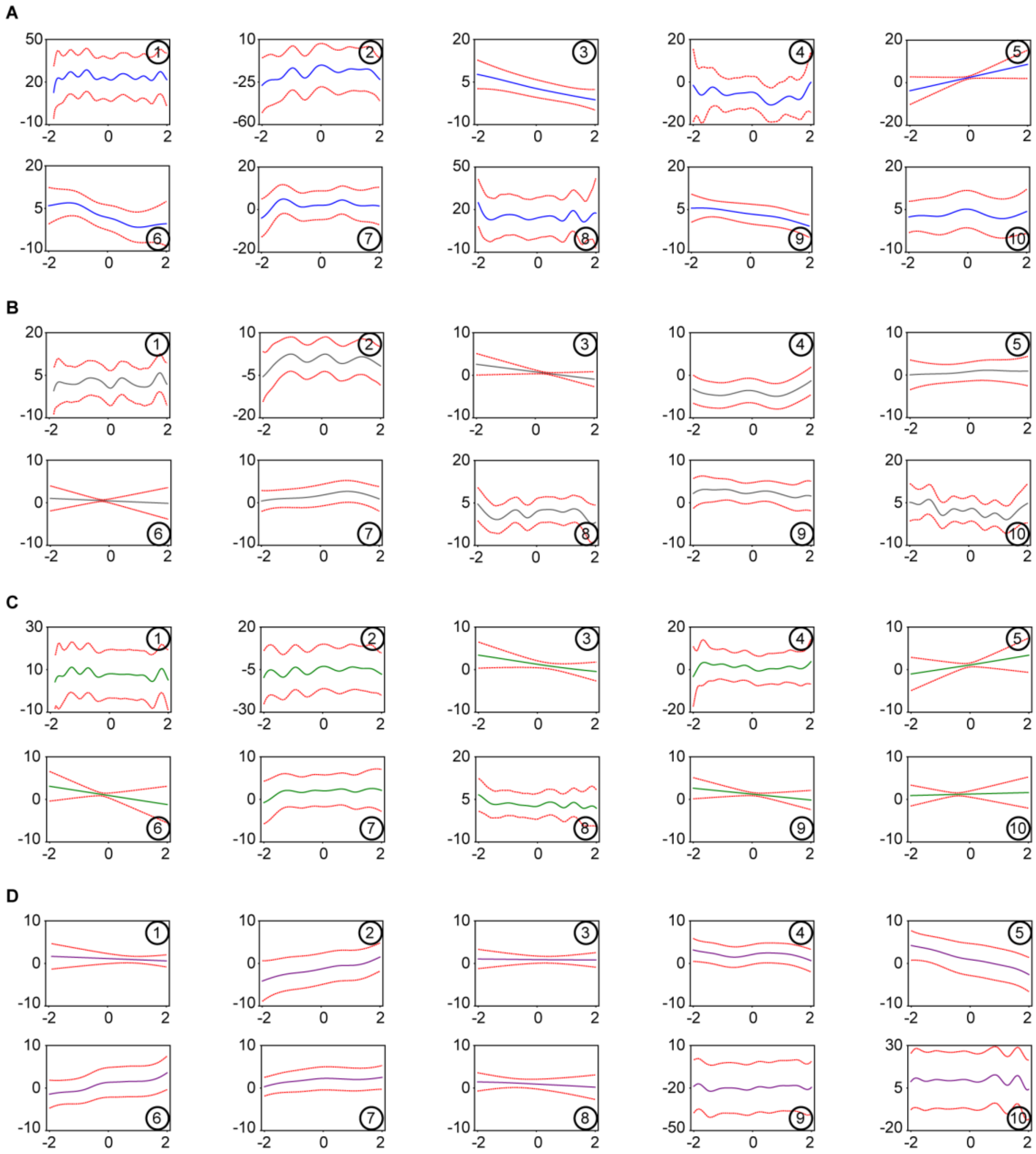


**Figure S6.** Partial dependence plots (PDPs) of gradient features when predicting PANSS total scores (**A**) and subscale scores of negative (**B**) and general psychopathology (**C**) as well as CFT scores (**D**). Features (clusters) were labelled with circle numbers.

**Table S1.** Demographics and clinical assessments for SZ and NC groups in the discovery and replication datasets.

| | Discovery dataset (n = 625) | | | | Replication dataset (n = 415) | | | |
|---|---|---|---|---|---|---|---|---|
| | SZ (n = 318) | NC (n = 307) | *P* value | N missing | SZ (n = 195) | NC (n = 220) | *P* value | N missing |
| **Demographics** | | | | | | | | |
| Age (y) | 27.1 ± 7.0 | 27.5 ± 6.5 | 0.48 | 0 | 27.9 ± 7.2 | 29.3 ± 7.7 | 0.07 | 0 |
| Gender (M / F) | 166 / 152 | 160 / 147 | 0.95 | 0 | 96 / 99 | 102 / 118 | 0.62 | 0 |
| **Clinical assessments** | | | | | | | | |
| PANSS total | 81.9 ± 11.6 | - | n/a | 0 | 86.9 ± 12.3 | - | n/a | |
| PANSS positive | 23.8 ± 4.1 | - | n/a | 0 | 24.4 ± 4.0 | - | n/a | 0 |
| PANSS negative | 19.3 ± 5.9 | - | n/a | 0 | 21.2 ± 5.7 | - | n/a | 0 |
| PANSS general | 38.8 ± 6.9 | - | n/a | 0 | 41.3 ± 6.9 | - | n/a | 0 |
| CFT | 15.7 ± 5.5 | - | n/a | 11 | 14.6 ± 5.1 | - | n/a | 5 |

PANSS, Positive and Negative Syndrome Scale; CFT, Category Fluency Test-Animal Naming; y, year; M, male; F, female; n/a = not applicable.

Data were presented as mean ± standard deviation.

**Table S2.** Significant clusters of differences in gradient between SZ and NC groups in the discovery dataset.

| Cluster | Cluster size | T value | Effect size | Brain region | MNI coordinate | | |
|---|---|---|---|---|---|---|---|
| | | | | | X | Y | Z |
| 1 | 57 | 4.53 | 0.41 | Frontal_Inf_Orb_L (aal) | -31 | 27 | -18 |
| 2 | 94 | 5.19 | 0.46 | Frontal_Inf_Orb_R (aal) | 43 | 40 | -7 |
| 3 | 34 | 5.07 | 0.45 | Temporal_Pole_Mid_L (aal) | -42 | 15 | -26 |
| 4 | 61 | 5.94 | 0.52 | Frontal_Inf_Tri_R (aal) | 51 | 22 | 21 |
| 5 | 64 | -4.08 | -0.38 | SupraMarginal_R (aal) | 60 | -22 | 26 |
| 6 | 34 | -3.88 | -0.36 | Insula_R (aal) | 38 | -6 | 13 |
| 7 | 52 | -4.31 | -0.39 | Parietal_Inf_L (aal) | -34 | -47 | 58 |
| 8 | 64 | -5.34 | -0.48 | Parietal_Sup_R (aal) | 25 | -61 | 49 |
| 9 | 37 | -4.35 | -0.40 | Precuneus_L (aal) | -11 | -63 | 59 |
| 10 | 58 | -4.29 | -0.39 | Parietal_Sup_L (aal) | -20 | -64 | 47 |

Cluster size was the number of vertices. Effect size was the score of Cohen's d.

**Table S3.** In the replication dataset, cluster-based replicated results of between-group differences in gradient.

| Cluster | T value | *P* value | FDR<0.05 | FDR<0.005 | FDR<0.001 |
|---|---|---|---|---|---|
| 1 | 3.80 | 1.6e-04 | T | T | T |
| 2 | 5.38 | 1.2e-07 | T | T | T |
| 3 | 3.47 | 5.8e-04 | T | T | F |
| 4 | 3.10 | 2.0e-03 | T | T | F |
| 5 | -2.52 | 0.011 | T | F | F |
| 6 | -2.04 | 0.035 | T | F | F |
| 7 | -0.40 | 0.68 | F | F | F |
| 8 | -1.95 | 0.042 | T | F | F |
| 9 | -4.27 | 2.4e-05 | T | T | T |
| 10 | -3.35 | 8.8e-04 | T | T | F |

T or F, survived or failed FDR correction at the significant level.

**Table S4.** In the discovery dataset, prediction of PANSS positive by linear generalized additive models (GAM).

| Outer fold | MAE | Pearson's r |
|---|---|---|
| 1 | 3.88 | 0.13 |
| 2 | 4.24 | 0.09 |
| 3 | 4.19 | 0.15 |
| 4 | 4.34 | -0.008 |
| 5 | 5.36 | 0.06 |
| Average | 4.40 | 0.08 |

Searching space for hyperparameters and training and testing procedures were identical with figure 3 in the main text.

**Table S5.** In the discovery dataset, prediction of clinical domains by other algorithms.

| Clinical domain | Algorithm | MAE | Pearson's r | *P* value |
|---|---|---|---|---|
| PANSS total | Linear regression | / | 0.06 | ns |
| | Lasso | / | 0.05 | ns |
| | Linear SVR | / | -0.01 | ns |
| | Kernel SVR | 12.42 | 0.14 | 0.018 |
| | Random Forest | 11.97 | 0.19 | 0.002 |
| PANSS positive | Linear regression | / | -0.04 | ns |
| | Lasso | / | -0.09 | ns |
| | Linear SVR | / | 0.02 | ns |
| | Kernel SVR | / | 0.10 | ns |
| | Random Forest | / | 0.02 | ns |
| PANSS negative | Linear regression | / | -0.02 | ns |
| | Lasso | / | 0.10 | ns |
| | Linear SVR | / | -0.03 | ns |
| | Kernel SVR | 7.41 | 0.17 | 0.004 |
| | Random Forest | / | 0.04 | ns |
| PANSS general | Linear regression | / | 0.09 | ns |
| | Lasso | / | 0.05 | ns |
| | Linear SVR | / | 0.03 | ns |
| | Kernel SVR | 7.72 | 0.16 | 0.008 |
| | Random Forest | 7.39 | 0.16 | 0.009 |
| CFT | Linear regression | / | 0.04 | ns |
| | Lasso | / | -0.1 | ns |
| | Linear SVR | / | 0.08 | ns |
| | Kernel SVR | / | 0.06 | ns |
| | Random Forest | / | 0.004 | ns |

Single 5-fold CV was applied in Linear regression, since it had no hyperparameters to tune. As for other models, nested 5-fold CV was performed. All other procedures and performance evaluation were the same with Figure 3 in the main text. ns, not significant. MAE was not shown (/) if ns.

Hyperparameters and search grids for each model were as follows: Lasso, alpha = $10^{-3}$, $10^{-2}$, $10^{-1}$, 0.2, 0.4, 0.6, 0.8. Linear SVR, C = $10^{-3}$, $10^{-2}$, $10^{-1}$, 1, 10, $10^{2}$, $10^{3}$. Kernel SVR, C = $10^{-3}$, $10^{-2}$, $10^{-1}$, 1, 10, $10^{2}$, $10^{3}$ and kernel = poly, rbf, sigmoid. Random Forest, n_estimators = 100, 200, 300, 400, 500, 600, 700, 800, 900, 1000 and max_features = auto, sqrt, log2.

**Table S6.** In the discovery dataset, variance inflation factor (VIF) was calculated to detect multicollinearity of features.

| Feature | 1 | 2 | 3 | 4 | 5 | 6 | 7 | 8 | 9 | 10 |
|---|---|---|---|---|---|---|---|---|---|---|
| VIF | 2.47 | 2.59 | 1.85 | 1.74 | 4.21 | 4.73 | 2.44 | 2.64 | 1.88 | 2.84 |

**Table S7.** In the discovery dataset, permutation feature importance was performed in regression analyses.

| Feature | PANSS total | | PANSS negative | | PANSS general | | CFT | |
|---|---|---|---|---|---|---|---|---|
| | r | Δr | r | Δr | r | Δr | r | Δr |
| 1 | 0.182 | 35% | 0.141 | 35.9% | 0.164 | 36.9% | 0.197 | 21.2% |
| 2 | 0.202 | 27.9% | 0.140 | 36.4% | 0.156 | 40% | 0.221 | 11.6% |
| 3 | 0.210 | 25% | 0.132 | 40% | 0.158 | 39.2% | 0.202 | 19.2% |
| 4 | 0.163 | 41.8% | 0.116 | 47.3% | 0.127 | 51.2% | 0.200 | 20% |
| 5 | 0.227 | 18.9% | 0.161 | 26.8% | 0.182 | 30% | 0.214 | 14.4% |
| 6 | 0.217 | 22.5% | 0.160 | 27.3% | 0.172 | 33.8% | 0.208 | 16.8% |
| 7 | 0.231 | 17.5% | 0.170 | 22.7% | 0.183 | 29.6% | 0.192 | 23.2% |
| 8 | 0.220 | 21.4% | 0.139 | 36.8% | 0.159 | 38.8% | 0.197 | 21.2% |
| 9 | 0.225 | 19.6% | 0.159 | 27.7% | 0.175 | 32.7% | 0.196 | 21.6% |
| 10 | 0.226 | 19.3% | 0.150 | 31.8% | 0.167 | 35.8% | 0.159 | 36.4% |

For each feature, r (Pearson correlation) was the median performance in 100 times of randomly shuffles. Δr was the decreased percentage of r relative to baseline score in figure 3 in the main text.
